# Comparative analysis of chloroplast genomes indicated different origin for Indian Tea (*Camellia assamica*) cv TV-1 as compared to Chinese tea

**DOI:** 10.1101/2020.07.13.199133

**Authors:** Hukam C. Rawal, Abhishek Mazumder, Sangeeta Borchetia, Biswajit Bera, S. Soundararajan, R Victor J Ilango, Anoop Kumar Barooah, Nagendra Kumar Singh, Tapan Kumar Mondal

## Abstract

Tea is an important plantation crop of some Asian and African countries. Based upon the morphological characteristics, tea is classified botanically into 2 main types i.e. Assam and China, which are morphologically very distinct. Further, they are so easily pollinated among themselves, that a third category, Cambod type is also described. Although the general consensus of origin of tea is India, Burma and China joining area, yet specific origin of China and Assam tea are not yet clear. In the present study, we made an attempt to understand the origin of Indian tea through the comparative analysis of different chloroplast (cp) genomes under the Camellia genus. Cp genome based phylogenetic analysis indicated that Indian Assam Tea, TV-1 formed a different group from that of China tea, indicating that TV-1 might have undergone different domestication and hence owe different origin. The simple sequence repeats (SSRs) analysis and codon usage distribution pattern also supported the clustering order in the cp genome based phylogenetic tree.

## 1. Introduction

Tea is a woody, perennial highly cross-pollinated crop so much so that the genus is very dynamic as indicated with the recent discovery of several new species (Lee and Yang 2019).More than 350 species are available today in this genus Camellia (Mukhopadhaya et al., 2016).Tea is such a natural drink which is taken by the majority of the world population in the morning.However among them, mainlytwo species i.e. *Camellia sinensis* (L.) O. Kuntz and *C. assamica* produce tea that we drink in various forms such as black tea, green tea or oolong tea (Mondal et al., 2004). However, in some pockets of China, tea is also made from a different species i.e. *C. taliensis*. Thus based upon the morphological characters, they are divided broadlyinto three types such as China tea, Assam tea and a hybrid between them called Cambod tea. Due to their high out-crossing nature, they breed freely among themselves which produce plant type that isintermediated between two extreme forms i.e. Assam type big leaf and China type small leaf. Thus,taxonomic classification of tea based on morphology is still confusing and species name is also a misnomer. For example, though China tea and Assam tea are considered to be separate species yet there is no restriction of gene flow. In factdesirable traits such as anthocyanin pigmentation or special quality characters of Darjeeling tea might have introduced from two wild species such as *C. irrawadiensis* and *C. taliensis* (Wood and Barua 1958). Besides taxonomy, origin of tea is debatable and needs to be clarified. Although Indo-Burma region near Irrawaddyriver is considered to be the centre of origin (Mondal 2009), yet due it is not clear weather Assam and China type tea have same or different domestication origin.

Chloroplasts (cp) are essential organelles that are primarily responsible for photosynthesis and hence found from green algae to higher plants (Xiong et al., 2009).They are maternally inherited and do not participate in genetic recombinationand hence are highly conserved (Palmer et al., 1988). The cp genome is circular with double stranded DNA molecule with a length of about 120–220 kb; that codes for 100-200 genes including protein-coding genes, rRNA and tRNA genes (Rogalski, et al., 2015). It also has a highly conserved quadripartite structure that includes, large single copy (LSC) region and a small single copy (SSC) region which is separated by two copies of inverted repeat regions (IR-A and IR-B). Due to this high conservation of chloroplast genomes compared to nuclear and mitochondrial genomes (Bi et al., 2018), they are widely used to differentiate closely related taxa, particularly some taxa below the species level with unclear taxonomic relationships. Complete chloroplast genome isthus used for deciphering phylogenetic relationships between closely related taxa to improve the understanding of the evolution of plant species. In the present study, we have used well annotated cp genome sequences of 17 species from Camellia genus including China type (*C. sinensis*), Assam type (*C. sinensis* var. assamica) and typical Indian Tea (Indian Assam type, TV1) that we decoded recently to understand the domestication, evolution of Indian tea and for exploiting DNA barcodes to geographically specific tea variety and for promoting germplasm innovation.

## 2. Materials and Methods

### 2.1. Data collection

To perform the chloroplast based phylogenetic analysis, we have chosen previously published and well-annotated cp genome sequences of 15 Camellia species (*Camellia sinensis*, *C. grandibracteata, C. leptophylla, C. petelotii, C. pubicosta, C. reticulata, C. azalea, C. japonica, C. chekiangoleosa, C. cuspidata, C. danzaiensis, C. impressinervis, C. pitardii, C. yunnanensis* and *C. taliensis*) (Huang et al., 2014) and 2 recently published cp genome sequences including *C. assamica* (Rawal et al., 2019) and *C. sinensis* var. *assamica* (Zhang et al., 2019). All these 17 Camellia species belongs to the Theaceae family under Asterids group. Moreover, in the analysis, while *Oryza sativa* and *Arabidopsis thaliana* were included as model monocot and dicot species, respectively, a bryophyte *Marchantia paleacea* was included to serve as the outgroup. We have also included 6 other dicot species comprising *Vitis vinifera* (grapes), *Solanum lycopersicum* (tomato) and *Coffea arabica* (coffee), which all three are believed to be evolutionary close to tea plant, 2 members from Asterids group but from family other than Theaceae i.e. *Actinidia tetramera* (crab-apple kiwi) and *Vaccinium macrocarpon* (cranberry), representing the family Actinidiaceae and Ericaceae, respectively. The cp genome sequences were downloaded for these 25 species from the NCBI Organelle Genome Resources database (Table 1).

**Table 1.**
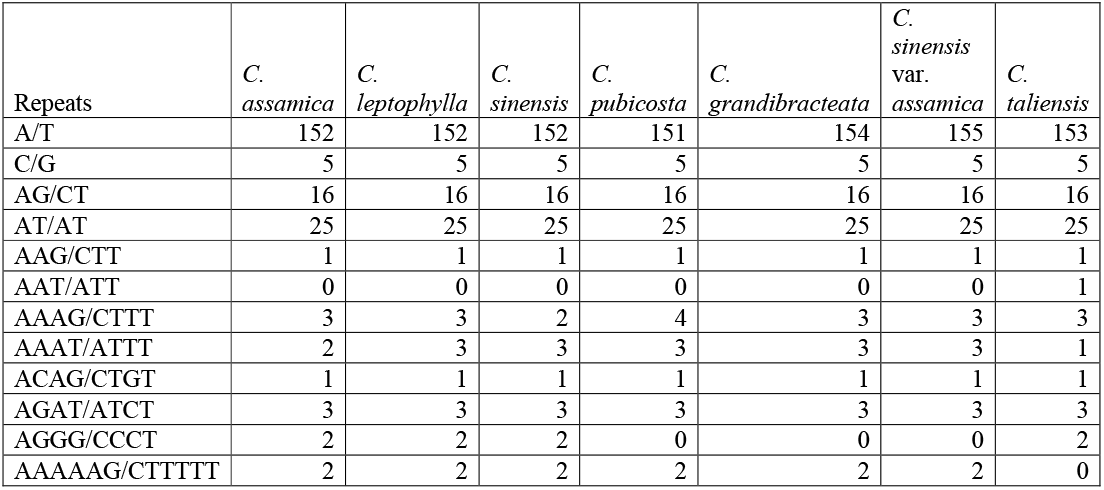
Frequency of classified SSR repeat types in 7 cp genomes

### 2.2. Phylogenetic analysis

We have used the method previously described in our previous study (Rawal et al., 2019). The cp genome sequences of all selected 25 species were aligned by MAFFT v7.402 (Katoh and Standley, 2013) at default parameters and the best-fit model for the downstream phylogenetic analysis was determined by ModelTest (ModelTest-NG v0.1.3) (Posada and Crandall, 1998). Finally, the Maximum likelihood (ML) tree was generated with RAxMLv8.2.12 (Stamatakis, 2014) by using the best-fit substitution model (GTRGAMMAX model) with 1000 bootstrap replicates. *M. paleacea*, a bryophyte served as the out-group in this analysis.

### 2.3. Comparative anallysis of seven Camellia species

Seven cp genomes of Camellia genus were further shortlisted based upon the results of phylogenetic analysis and compared for their cp genomic features including length, LSC and SSC and IR regions number of protein-coding genes, tRNA and rRNA genes. Further simple sequence repeats (SSRs) were identified in these cp genomes by MIcroSAtellite identification tool (MISA) (Thiel et al., 2003) with a threshold of 8, 4, 4, 3, 3 and 3 for minimum number of nucleotide repeats in mono-, di-, tri-, tetra-, penta- and hexa-nucleotide repeats, respectively.

For a specified codon, the Relative Synonymous Codon Usage (RSCU) value is the ratio of its actual usage frequency to expected frequency when it is used without bias. These RSCU values were calculated for seven Camellia species using ACUA (Vetrivel et al., 2007) for 60 codons (excluding 3 stop and 1 start codon from the total 64 codons).

## 3. Results and discussion

### 3.1. Phylogenetic analysis based on cp genomes

The ML tree based on alignment of cp genome sequences using GTRGAMMAX model at 1000 bootstrap replicates supported the branching-off of the bryophyte *M. paleacea*, an outgroup in this analysis, from rest of the plant species and serving as the root for the tree. *O. sativa*, the only monocot included in the study, is the next to branched off from rest of the species (dicots) in the tree (Fig. 1). *V. vinifera* and *A. thaliana* (the two Rosids member) are the next species to get separated from rest of the species, all of which belongs to Asterids group. Among Asterids, *C. Arabica* and *S. lycopersicum* are the first one to get separated from the remaining species under analysis, all of which represents the order Ericales. These species under Ericales includes *A. tetramera, V. macrocarpon* and all Camellia species, representing three families including Actinidiaceae, Ericaceae and Theaceae, respectively.

**Fig. 1.**
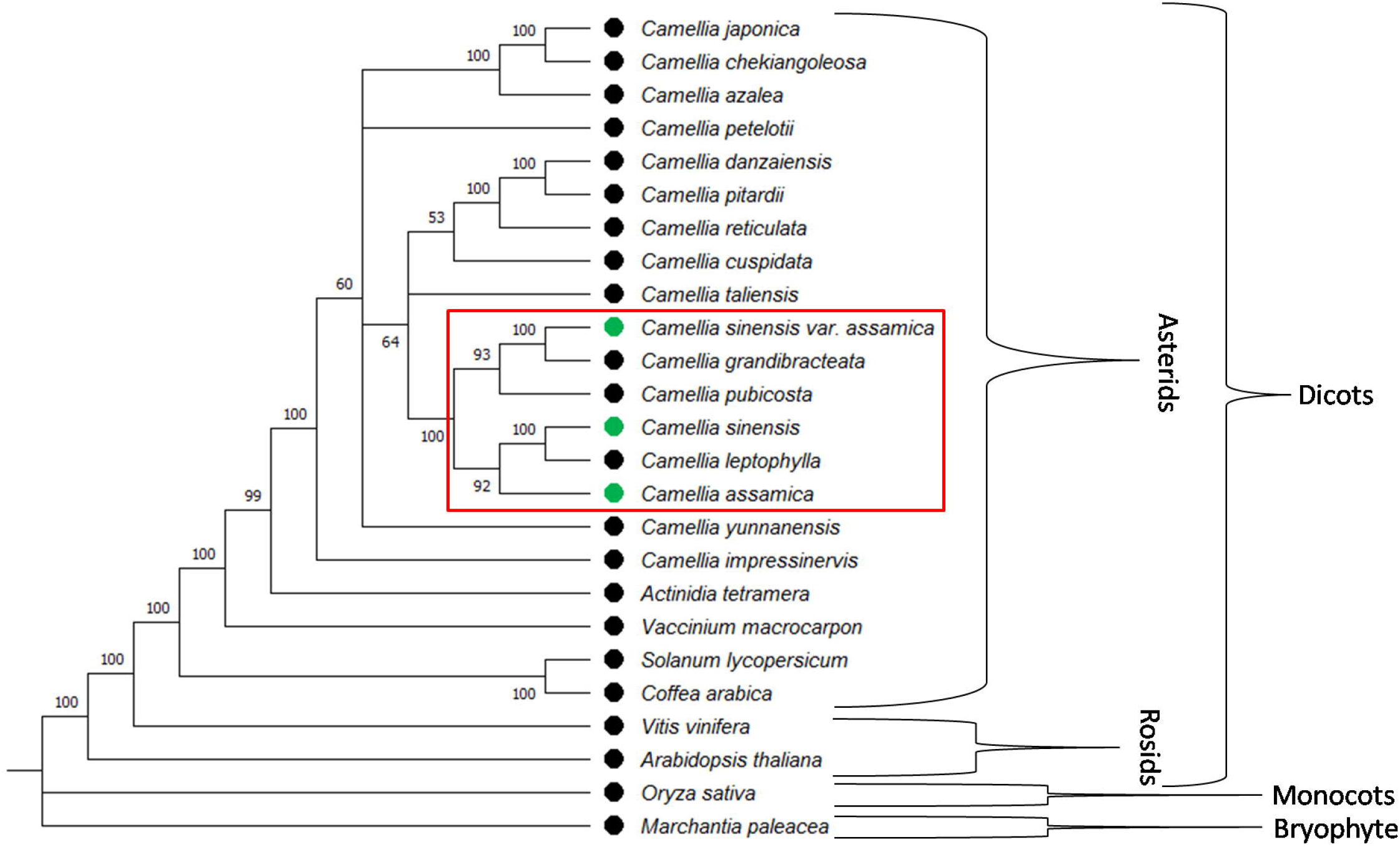
Chloroplast (cp) genome sequence based ML tree showing the phylogenetic relation between China tea, China Assam tea and Indian Assam tea (green bullet) to confirm their different origins

Among Camellia genus or Theaceae family, all three domesticated tea *C. assamica* (Indian Assam tea), *C. sinensis* and *C. sinensis* var. *assamica* (China Assam tea) are present in a single major cluster adjacent to the single clade of *C. taliensis*, a well known close wild relative of domesticated tea (Lie et al., 2015). Here within this cluster, we observed that Indian Assam tea and China Assam tea belongs to different clades and must have different origins as predicted previously (Wambulwa et al., 2016 and Meegahakumbura et al., 2016). Interestingly, *C. sinensis* was found evolutionary close to Indian Assam tea and seems to be branched-off from Indian Assam tea along with *C. leptophylla* in a similar manner and time as the China Assam tea and *C. grandibracteata* gets branched-off from *C. pubicosta*. This presence of *C. leptophylla* as sister clade with *C. sinensis* and existence of *C. pubicosta* as sister clade to *C. sinensis* var. *assamica* and *C. grandibracteata* is consistent with the previous study (Huang et al., 2014). Moreover, both China type tea (*C. sinensis* and China Assam tea) seems to be evolved after Indian Assam tea. Similar phylogenetic relations were found among these seven Camellia species (*C. assamica*, *C. leptophylla*, *C. sinensis*, *C. pubicosta, C. grandibracteata*, *C. sinensis* var. *assamica*, and *C. taliensis*) when inferred with other different and fast phylogenetic tree construction methods in MEGAX (Kumar et al., 2018) including Unweighted Pair Group Method with Arithmetic mean (UPGMA) (Fig. S1), Neighbour-joining (NJ) and Minimum Evolution method with strong bootstrap supports.

### 3.2. Comparative study of seven Camellia species

A comparative study of cp genome features of these seven species revealed that there is hardly any differences or patterns to notice in terms of cp genome size, LSC, SSC or IR region, number of genes, tRNAs and rRNAs (Table S2) to correlate with their phylogenetic positions in the tree. Even list of all genes found almost similar and conserved and has no clues about their relations (Table S3). Interestingly, distribution and frequency of SSR repeat types follow the pattern that these seven genomes have in the phylogenetic tree(Table S4 and S5). Distribution of SSR types shows that *C. taliensis*, a wild relative to domesticated tea, has a different pattern with 2 tri-mers as compared to 1 in other six cp genomes and no hexa-mers as compared to 2 in other six (Table 1, S4 and Fig. 2 a). No AAT/ATT motif was found in these cp genomes, except 1 in *C. taliensis* and two AAAAAG/CTTTTT motifs were there in these six genomes but none in *C. taliensis* (Table 1, Fig. 2b). More importantly, AGGG/CCCT motif has frequency of two in each of *C. assamica*, *C. leptophylla* and *C. sinensis* (which are present in one cluster in the tree) but totally missing in *C. pubicosta, C. grandibracteata* and *C. sinensis* var. *assamica* (which altogether represents a different cluster in the tree).

**Fig. 2.**
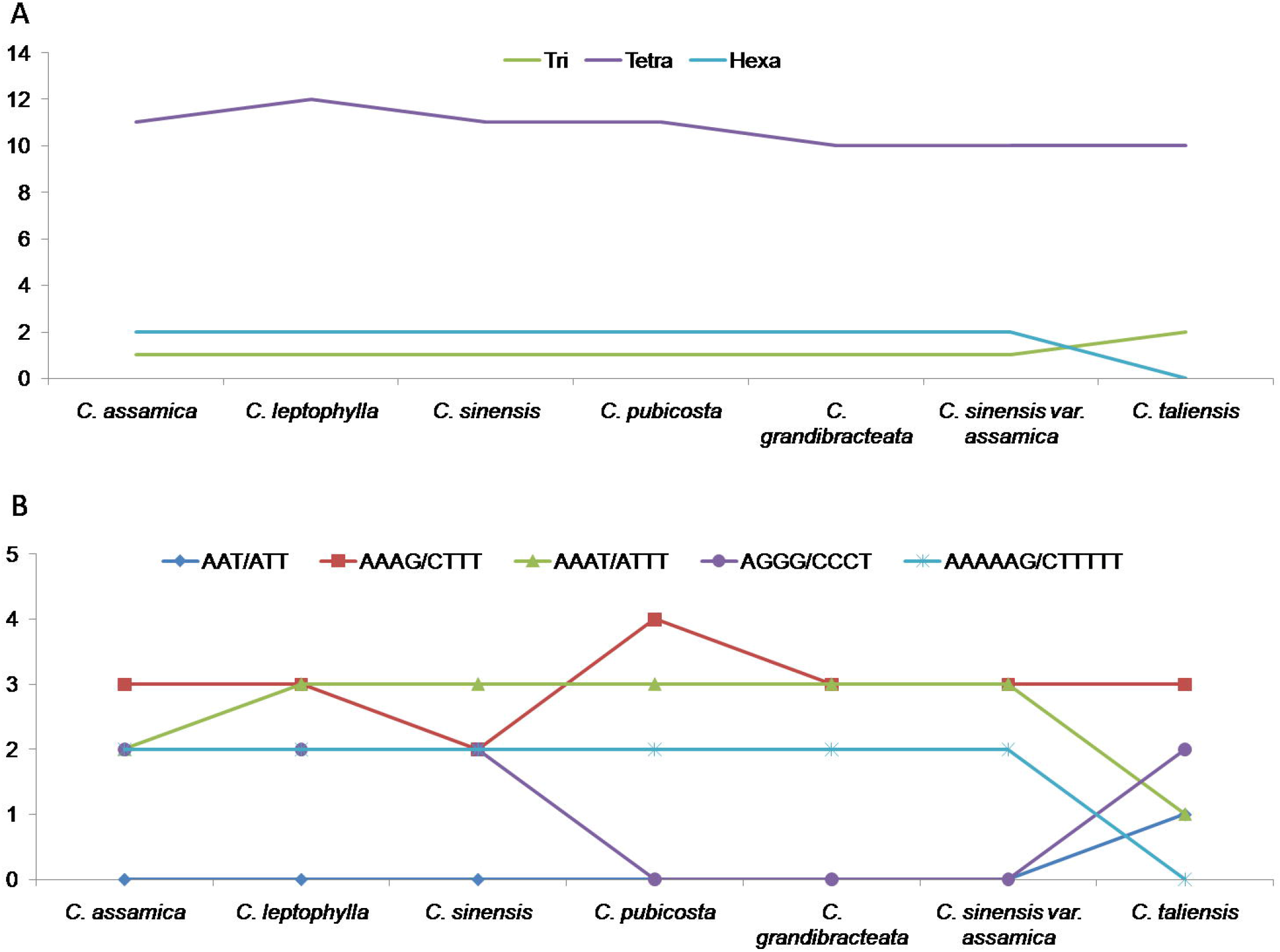
Distribution pattern of tri, tetra and hexa type SSR number (A) and SSR motifs (B) showing related variation between first three (*C. assamica*, *C. leptophylla* and *C. sinensis*), next three (*C. pubicosta, C. grandibracteata* and *C. sinensis* var. *assamica*) or last species (*C. taliensis*) under analysis

The codon usage frequency distribution among seven Camellia species showed that Indian Assam type tea has a different pattern than the rest of the six Chinese cp genomes (Table S6). Among these, when we further compared this codon usage frequency of Indian tea with two other domesticated tea (*C. sinensis* and *C. sinensis* var. *assamica*) and one of their close wild relative *C. taliensis*, we observed that out of 60 codons, only 5 codons have similar RSCU values in all 4 cp genomes (Table 2). Among rest of the 55 codons, the distribution pattern of 39 codons (marked with # sign) in Indian tea was found different from other three Chinese tea and only 7 were similar with *C. sinensis* and 6 were similar with each of *C. sinensis* var. *assamica* and *C. taliensis*. Hence, out of total 60 codons, 65% codons have different patterns in Indian tea, 11% shared with *C. sinensis*, 10% shared with each of *C. sinensis* var. *assamica* and *C. taliensis* and around 8% shared common pattern between all four. Apart from similar in all 4 cp genomes, the three Chinese genomes shared a very good number of similar codon usage patterns with 23 between *C. sinensis* and *C. sinensis* var. *assamica*, 30 between *C. sinensis* var. *assamica* and *C. taliensis*, and 25 between *C. sinensis* and *C. taliensis*. Overall, these results indicated that while the Indian cp genome has different codon usage patterns, the three Chinese cp genomes have quite similar patterns. Hence there must be different codon usage selection between cp genomes of Indian and Chinese tea with possible chances of different domestications and origins of Indian and Chinese tea.

**Table 2.** The codon usage frequency distribution in terms of RSCU values among four cp genomes of tea with colour coded for blue-yellow-red scale for maximum to minimum values. With different remarks for identical RSCU values in all 4 cp genomes (*), between *C. sinensis* var. *assamica* and *C. sinensis* (CSA-CSS), between *C. sinensis* and *C. taliensis* (CSS-CST), between *C. sinensis* var. *assamica* and *C. taliensis*(CSA-CST), between C*. assamica* and *C. sinensis* (Ind-CSS), between *C*. *assamica* and *C*. var. *assamica* (Ind-CSA), between *C. assamica* and *C. taliensis* (Ind-CST), between all three Chinese cp genomes (CSA-CSS-CST) and different RSCU values in *C. assamica* as compared to three Chinese cp genomes (#)

| Amino Acid | Codon | *C. assamica* | *C. sinensis* | *C. sinensis* var. *assamica* | *C. taliensis* | Remarks |
| --- | --- | --- | --- | --- | --- | --- |
| A | GCA | 1.11 | 1.14 | 1.14 | 1.15 | #,CSA-CSS |
| A | GCC | 0.66 | 0.65 | 0.64 | 0.65 | #,CSS-CST |
| A | GCG | 0.39 | 0.39 | 0.38 | 0.38 | Ind-CSS,CSA-CST |
| A | GCT | 1.83 | 1.83 | 1.84 | 1.83 | Ind-CSS,Ind-CST,CSS-CST |
| C | TGC | 0.49 | 0.47 | 0.48 | 0.47 | #,CSS-CST |
| C | TGT | 1.51 | 1.53 | 1.53 | 1.53 | #,CSA-CSS-CST |
| D | GAC | 0.38 | 0.37 | 0.37 | 0.36 | #,CSA-CSS |
| D | GAT | 1.62 | 1.63 | 1.63 | 1.64 | #,CSA-CSS |
| E | GAA | 1.49 | 1.51 | 1.51 | 1.51 | #,CSA-CSS-CST |
| E | GAG | 0.51 | 0.49 | 0.49 | 0.49 | #,CSA-CSS-CST |
| F | TTC | 0.73 | 0.71 | 0.72 | 0.72 | #,CSA-CST |
| F | TTT | 1.27 | 1.29 | 1.28 | 1.28 | #,CSA-CST |
| G | GGA | 1.63 | 1.64 | 1.64 | 1.65 | #,CSA-CSS |
| G | GGC | 0.42 | 0.42 | 0.41 | 0.41 | Ind-CSS,CSA-CST |
| G | GGG | 0.68 | 0.68 | 0.67 | 0.67 | Ind-CSS,CSA-CST |
| G | GGT | 1.28 | 1.27 | 1.28 | 1.27 | Ind-CSA,CSS-CST |
| H | CAC | 0.44 | 0.43 | 0.43 | 0.43 | #,CSA-CSS-CST |
| H | CAT | 1.56 | 1.58 | 1.57 | 1.57 | #,CSA-CST |
| I | ATA | 0.93 | 0.96 | 0.96 | 0.96 | #,CSA-CSS-CST |
| I | ATC | 0.61 | 0.60 | 0.60 | 0.60 | #,CSA-CSS-CST |
| I | ATT | 1.46 | 1.44 | 1.44 | 1.44 | #,CSA-CSS-CST |
| K | AAA | 1.46 | 1.49 | 1.48 | 1.49 | #,CSS-CST |
| K | AAG | 0.54 | 0.51 | 0.52 | 0.51 | #,CSS-CST |
| L | CTA | 0.81 | 0.80 | 0.81 | 0.81 | Ind-CSA,Ind-CST,CSA-CST |
| L | CTC | 0.46 | 0.43 | 0.45 | 0.43 | #,CSS-CST |
| L | CTG | 0.39 | 0.39 | 0.39 | 0.39 | * |
| L | CTT | 1.25 | 1.25 | 1.26 | 1.27 | Ind-CSS |
| L | TTA | 1.85 | 1.91 | 1.89 | 1.90 | # |
| L | TTG | 1.24 | 1.22 | 1.20 | 1.20 | #,CSA-CST |
| N | AAC | 0.47 | 0.46 | 0.46 | 0.46 | #,CSA-CSS-CST |
| N | AAT | 1.53 | 1.54 | 1.54 | 1.54 | #,CSA-CSS-CST |
| P | CCA | 1.18 | 1.19 | 1.18 | 1.19 | Ind-CSA,CSS-CST |
| P | CCC | 0.70 | 0.70 | 0.70 | 0.70 | * |
| P | CCG | 0.50 | 0.50 | 0.51 | 0.51 | Ind-CSS,CSA-CST |
| P | CCT | 1.63 | 1.60 | 1.61 | 1.61 | #,CSA-CST |
| Q | CAA | 1.51 | 1.53 | 1.53 | 1.53 | #,CSA-CSS-CST |
| Q | CAG | 0.49 | 0.48 | 0.47 | 0.47 | #,CSA-CST |
| R | AGA | 1.78 | 1.86 | 1.86 | 1.86 | #,CSA-CSS-CST |
| R | AGG | 0.63 | 0.62 | 0.62 | 0.61 | #,CSA-CSS |
| R | CGA | 1.45 | 1.44 | 1.44 | 1.43 | #,CSA-CSS |
| R | CGC | 0.34 | 0.33 | 0.33 | 0.32 | #,CSA-CSS |
| R | CGG | 0.43 | 0.42 | 0.42 | 0.42 | #,CSA-CSS-CST |
| R | CGT | 1.37 | 1.33 | 1.32 | 1.35 | # |
| S | AGC | 0.35 | 0.34 | 0.34 | 0.34 | #,CSA-CSS-CST |
| S | AGT | 1.25 | 1.22 | 1.23 | 1.23 | #,CSA-CST |
| S | TCA | 1.14 | 1.19 | 1.19 | 1.19 | #,CSA-CSS-CST |
| S | TCC | 0.96 | 0.94 | 0.94 | 0.95 | #,CSA-CSS |
| S | TCG | 0.52 | 0.51 | 0.52 | 0.51 | Ind-CSA,CSS-CST |
| S | TCT | 1.79 | 1.80 | 1.78 | 1.79 | Ind-CST |
| T | ACA | 1.21 | 1.23 | 1.24 | 1.22 | # |
| T | ACC | 0.73 | 0.73 | 0.73 | 0.73 | * |
| T | ACG | 0.42 | 0.41 | 0.42 | 0.42 | Ind-CSA,Ind-CST,CSA-CST |
| T | ACT | 1.64 | 1.63 | 1.61 | 1.64 | Ind-CST |
| V | GTA | 1.49 | 1.50 | 1.49 | 1.50 | Ind-CSA,CSS-CST |
| V | GTC | 0.47 | 0.46 | 0.48 | 0.47 | Ind-CST |
| V | GTG | 0.56 | 0.56 | 0.55 | 0.55 | Ind-CSS,CSA-CST |
| V | GTT | 1.48 | 1.48 | 1.48 | 1.48 | * |
| W | TGG | 1.00 | 1.00 | 1.00 | 1.00 | * |
| Y | TAC | 0.38 | 0.40 | 0.39 | 0.39 | #,CSA-CST |
| Y | TAT | 1.62 | 1.61 | 1.61 | 1.61 | #,CSA-CSS-CST |

## Conclusion

In the present study, no deviation was observed from the known taxonomic classification from sub-class or order level to family or genus level by following this cp genome based phylogenetic relationship study. Thus one can rely on cp based evolutionary study and the observed phylogenetic relations among species under consideration. This analysis supported the possible different domestication origins of Indian Assam tea and China Assam tea with the existence of Indian Assam tea prior to *C. sinensis* and China Assam tea which is well supported with SSR and codon usage frequency distribution.

---

Table S1. List of chloroplast (cp) genome sequence used in the analysis

Table S2. Basic features in shortlisted 7 cp genomes

Table S3. List of genes in shortlisted 7 cp genomes

Table S4. Distribution to different repeat type classes

Table S5. Frequency of identified SSR motifs

Table S6. The codon usage frequency distribution in terms of RSCU values among seven cp genomes Camellia genus

Supplementary Fig. S1. UPGMA method based phylogenetic tree between 7 cp genome sequences of Camellia species. Sequences were aligned with ClustalW and Evolutionary analyses were conducted in MEGA X at 1000 bootstrap support.

