## Supplementary figures and images for "Comparative analysis of chloroplast genomes indicated different origin for Indian Tea (*Camellia assamica*) cv TV-1 as compared to Chinese tea"

### Supplementary Fig. S1

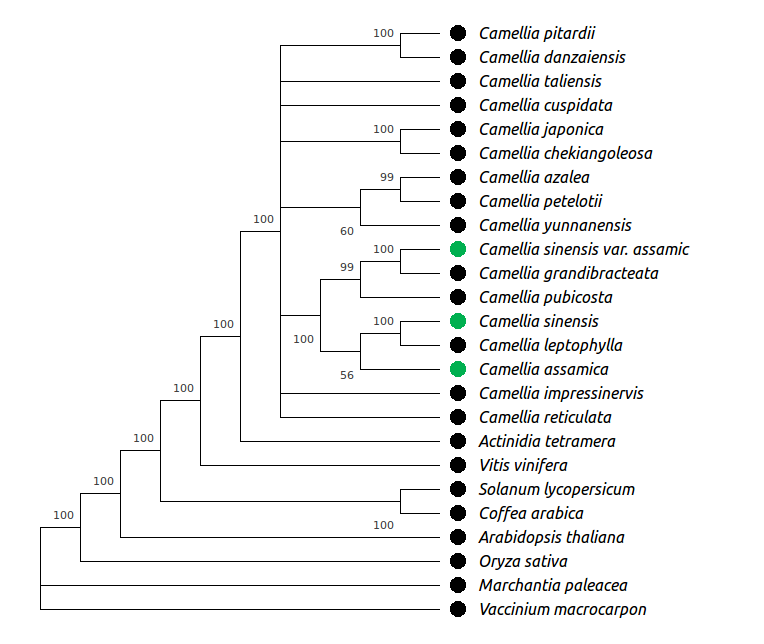
